# Habitat loss is information loss: Species distribution models are compromised in anthropogenic landscapes

**DOI:** 10.1101/258038

**Authors:** Russell Dinnage, Marcel Cardillo

## Abstract

Species distribution models (SDMs) are valuable tools to estimate species’ distributions, but are vulnerable to biases in the probability of a species being observed. One such bias is habitat loss, which has affected a substantial and increasing proportion of the Earth. In regions of severe habitat loss, data on a species’ occurrence may represent a small, non-random subset of sites it once occupied. This could cause distorted reconstructions of species distributions, and misleading inferences of evolutionary and ecological processes. We present a statistical approach for quantifying the influence on SDMs of habitat loss, and generating distribution predictions that are robust to these biases. We explored some of the effects of accounting for habitat loss on inferences from common downstream biogeographic and ecological analysis methods.

We used herbarium record data to model the distribution of 325 plant species in the genera *Banksia* and *Hakea* across Australia, using point process models. We accounted for biases in the models by including a proxy variable representing habitat loss, and compared the fit of models without this variable to those with it. We explored the influence of habitat loss by mapping biodiversity patterns predicted with and without accounting for it.

Generally, accounting for habitat loss in SDMs led to increases in the mean area of modelled species distributions of ~10% for *Banksia* and ~12% for *Hakea* across Australia (in some cases, up to several 100,000 km^2^ increases in predicted range), with somewhat greater average increases (11% and 15%) for species in the southwest Australian biodiversity hotspot. Accounting for habitat loss leads to an increase in predicted species richness (Alpha and Gamma diversity), but a decrease in compositional turnover (Beta diversity), across most of Australia. Accounting for habitat loss in SDMs had minimal influence on a downstream macroevolutionary analysis (Age-Range Correlation) that utilizes species distributions, seemingly because exposure to habitat loss did not show a phylogenetic pattern in this taxonomic group.

The influence of habitat loss on species distributions estimated with SDM is likely to be context-dependent and difficult to generalize, but will tend to cause underestimates of range sizes. This may have consequences for mapping spatial patterns of diversity and for some downstream analyses of biogeographic, evolutionary, or ecological processes, based on species distributions, as well as conservation measures that rely on accurate species mapping.

## Introduction

The description and mapping of species geographic distributions underpins many areas of research in ecology, evolution, and biogeography. For example, estimates of overlap in species distributions are used to infer the geographic mode of speciation (e.g. Anacker and Strauss 2014, Cardillo and Warren 2016), and infer the strength of interspecific competition (e.g. Davies et al. 2007); present-day distributions mapped onto phylogenies are used to infer ancestral distributions and reconstruct the biogeographic history of clades (Matzke 2013, Quintero et al. 2015); and the geographic patterns of species diversity or turnover in community composition are calculated from the conjunction of species distributions in different areas (Orme et al. 2005, Buckley and Jetz 2008). However, species distributions are rarely measured directly: usually, they are reconstructed from sets of occurrence records from specific locations, and the completeness and coverage of species records is often determined by practical constraints. For this reason, Species Distribution Modelling (SDM), a set of statistical methods that predict species distributions from their associations with environmental features, has emerged as a key tool in modern biodiversity research.

Although SDM algorithms have become quite sophisticated, they still can only extrapolate from observed species occurrences, which means they are vulnerable to biases arising from factors that influence the probability of an occurrence being measured. Sampling biases resulting from differential sampling intensity across a region (e.g. associated with access such as road networks) have received a lot of attention in the SDM literature (Elith and Leathwick 2007, Phillips et al. 2009, Syfert et al. 2013, Warton et al. 2013, Fernández and Nakamura 2015, Fithian et al. 2015, Guillera-Arroita 2016). Other kinds of bias, including habitat loss, have received remarkably little attention but may have a direct role in obscuring the patterns of species distribution from which we try to infer evolutionary or ecological processes, or plan conservation actions. In regions where much of the natural habitat has been destroyed or heavily modified, we expect to find fewer occurrence records for many species, especially since many such regions are unlikely to have been thoroughly sampled before the onset of habitat conversion. With greater than 50% of the world‘s natural habitat now converted to anthropogenic landscapes, and increasingly large, contiguous regions now almost entirely devoid of natural habitat (Ellis 2014), bias in occurrence records arising from habitat loss could have a potentially severe impact on SDM.

The anthropogenic distortion of species distributions is likely to be especially severe in biodiversity hotspots. With high levels of endemic plant species richness and turnover, hotspots have been identified as critical conservation focus regions (Myers et al. 2000). The other defining criterion of biodiversity hotspots is extensive habitat loss (Myers et al 2000). This makes hotspots, by definition, potentially susceptible to inaccurate reconstructions of species distributions, because habitat loss removes critical information needed for accurate predictions under SDM. This creates a double indemnity, whereby we suffer the future cost of losing species and populations themselves, as well as the cost of losing the valuable information their observation entails. This can then compound further, since it is often the information we infer about a species’ distribution, and their response to the environment implicit within it, that helps us to effectively conserve those species. It is therefore important to quantify the influence that habitat loss has on inferred species distributions, and develop methods to account for this using only the data on post-habitat loss species occurrences that are typically available.

Here, we present a methodological framework based on poisson point-process SDM modelling (hereafter referred to as PPM) to quantify and adjust for the influence of anthropogenic habitat loss on species distributions inferred with SDM. As case studies we use two species-rich plant genera of the Family Proteaceae (*Banksia* and *Hakea*) that are distributed widely across Australia, both within and outside the southwest Australian biodiversity hotspot. Our approach is to explicitly incorporate the degree of habitat loss into SDMs for *Banksia* and *Hakea* species, and to compare the fit of these models to models that do not consider habitat loss, as well as to models that incorporate the more well-studied sampling bias associated with road access. We infer species distributions under SDMs accounting for habitat loss, and investigate the extent to which distributions are over or under-predicted when habitat loss is ignored, with particular focus on the southwest biodiversity hotspot. We then investigate the influence of accounting for habitat loss in SDMs on reconstructions of continent-wide patterns of species diversity and turnover. Finally, we present an example of how accounting for habitat loss in SDMs can influence the results of a downstream macroevolutionary analysis that utilizes data on inferred species distributions.

## Methods

### Species occurrence and environmental data

We downloaded occurrence data for every species of *Banksia* and *Hakea* (173 and 152 species, respectively) from the Atlas of Living Australia (http://www.ala.org.au/). We cleaned the occurrence records dataset by removing records that were obviously incorrect (e.g. in the sea) or well outside the bounds of expert range maps (e.g. cultivated specimens in botanic gardens). We downloaded data for a large number of environmental variables from the Atlas of Living Australia, using the R (R Core Team 2016) package ALA4R (Raymond et al. 2015). This included 104 variables relating to climate, soil, and topographic characteristics (Table S1) for coordinates corresponding to all species occurrence records, as well as for ~3 million quadrature points (see below).

In order to model the distribution of species when only presence data are recorded (i.e. from occurrence records), we generated a set of comparison points, known as “quadrature points” in point process SDMs (Renner et al. 2015). We generated one point for each 0.015 degree of latitude and longitude across the whole of Australia (~3 million points in total). To apply PPM to species, we sampled quadrature points from all of the terrestrial ecoregions (Olson et al. 2001) in which at least one occurrence record for the species was found (Barve et al. 2011).

### Habitat Loss Proxy Variable

In order to statistically account for habitat loss in our occurrence data, we collected a variable representing modern habitat loss across Australia. In agricultural landscapes, much of the native vegetation has been completely or largely removed for agricultural development. For many species, there will be a greatly reduced probability that occurrence records from such areas exist, except where remnant native vegetation remains in patches large and numerous enough to support substantial populations. Even where the basic structure of the native vegetation is largely maintained (e.g. in some regions dominated by pastoral land uses), the habitat value of the vegetation may be severely diminished and species occurrence records few or non-existent. To quantify habitat loss we used data on dynamic vegetation land cover from Geoscience Australia (http://www.ga.gov.au/scientific-topics/earth-obs/landcover). This dataset uses satellite reflectance data to classify each 250x250m square of Australia into a set of vegetation classes (12 natural and 8 non-natural, including agricultural). For use in the models we converted this land cover dataset into a binary classification (1=natural, 0=non-natural) and then used this to calculate the distance from each occurrence point to the nearest natural area (See Supplementary Methods 1).

### Accounting for Sampling Bias

Many studies have shown that the distribution of species occurrence records is often closely associated with road networks, because roads provide easy access and the time and cost of collecting or surveying increases with distance from roads (Elith and Leathwick 2007, Phillips et al. 2009, Syfert et al. 2013, Warton et al. 2013, Fernández and Nakamura 2015, Fithian et al. 2015, Guillera-Arroita 2016). We wanted to account for this potential bias while examining the effects of habitat loss. This would be particularly important if habitat loss was highly correlated with road access (though this correlation was not as strong as suspected: see Supplementary Figure 5). For each occurrence and quadrature point, we calculated the distance to the nearest road using road data in OpenStreetMap (https://www.openstreetmap.org; See supplement for more details.) and used it to account for sampling bias following the procedure of (Warton et al. 2013).

### Variable selection and implementation of Point Process Models

Point Process Models (PPMs) are a class of Species Distribution models able to incorporate linear and polynomial terms of any included predictor (i.e. environmental factor). Given the large number of environmental predictors available from ALA, and no strongly-supported *a priori* framework to choose the predictors, we used a guided regularized random forest (GRRF) approach (Deng and Runger 2013) to conduct initial variable selection for each species, such that we only used the best eight variables in this analysis (see Supplementary Methods 2).

We used the ppmlasso package in R (Renner and Warton 2013) to fit our PPMs. The ppmlasso function also includes a lasso penalty algorithm to reduce the number of variables in the model to minimize the risk of over-fitting. Including a proxy variable to account for bias has a rich history in the analysis of survey data in the social sciences, where it has been shown that the relationship between a bias-related variable and the probability of inclusion in a dataset is often non-linear and interacts with predictors (Gelman 2007). On the assumption that this principle is likely to apply to biological record data as well, we allowed habitat loss variables to have a more complicated relationship to the probability of occurrence than has been done before (e.g. Warton et al. 2013). For each *Banksia* and *Hakea* species we fitted seven different models, allowing different levels of complexity concerning habitat loss (and sampling bias). The formulation of the models is summarized below.

*Model 1 (No Habitat Loss)*: Eight top environmental variables (8 terms) + all polynomial combinations of environmental variables of degree 2 (36 terms) (i.e. all environmental squared terms plus all pairwise interactions). Total terms: 44

*Model 2 (No Habitat Loss accounting for road distance)*: Model 1 (44 terms) + road distance (1 term) + road distance squared term (1 term). Total terms: 46

*Model 3 (Habitat Loss only)*: Model 1 (44 terms) + habitat loss (1 term) + habitat loss squared term (1 term). Total terms: 46

*Model 4 (Habitat Loss accounting for road distance)*: Model 1 (44 terms) + habitat loss and road distance (2 terms) + all polynomial combinations of habitat loss and road distance of degree 2 (3 terms). Total terms: 49

*Model 5 (Habitat Loss only + interactions)*: Model 1 (44 terms) + habitat hoss (1 term) + habitat loss squared term (1 term) + all polynomial combinations of environmental variables and habitat loss (8 terms). Total terms: 54

*Model 6 (No Habitat Loss accounting for road distance + interactions*: Model 1 (44 terms) + road distance (1 term) + road distance squared term (1 term) + all polynomial combinations of environmental variables and road distance (8 terms). Total terms: 54

*Model 7 (Full Model, or Habitat Loss accounting for Sampling Bias + interactions)*: Model 1 (44 terms) + habitat loss and road distance (2 terms) + all polynomial combinations of environmental variables, habitat loss and road distance (19 terms). Total terms: 65

Of these, variables with low explanatory power had their coefficients shrunk towards zero by a L1 lasso regularization path algorithm. Briefly, 20 models were fit, each with an increasing shrinkage parameter applied to the variables, such that increasingly more coefficients with low values would be set to zero. Out of this 20, the model with the lowest Bayesian Information Criterion (BIC) score was retained.

For each genus (*Banksia* and *Hakea*) we first tested whether including habitat loss proxy variables improved the overall predictive power of the models. We used the Akaike weights (Burnham and Anderson 2003) to test the relative fit of models containing different habitat loss versus those that did not (Figure 1). Akaike weights measure the strength of evidence favouring a predictive model over all other models fit to the data. Higher values correspond to a higher probability that the model was the best predictive model out of those tried. We averaged Akaike weights across all species to test the overall predictive performance of the seven models.

**Figure 1.**
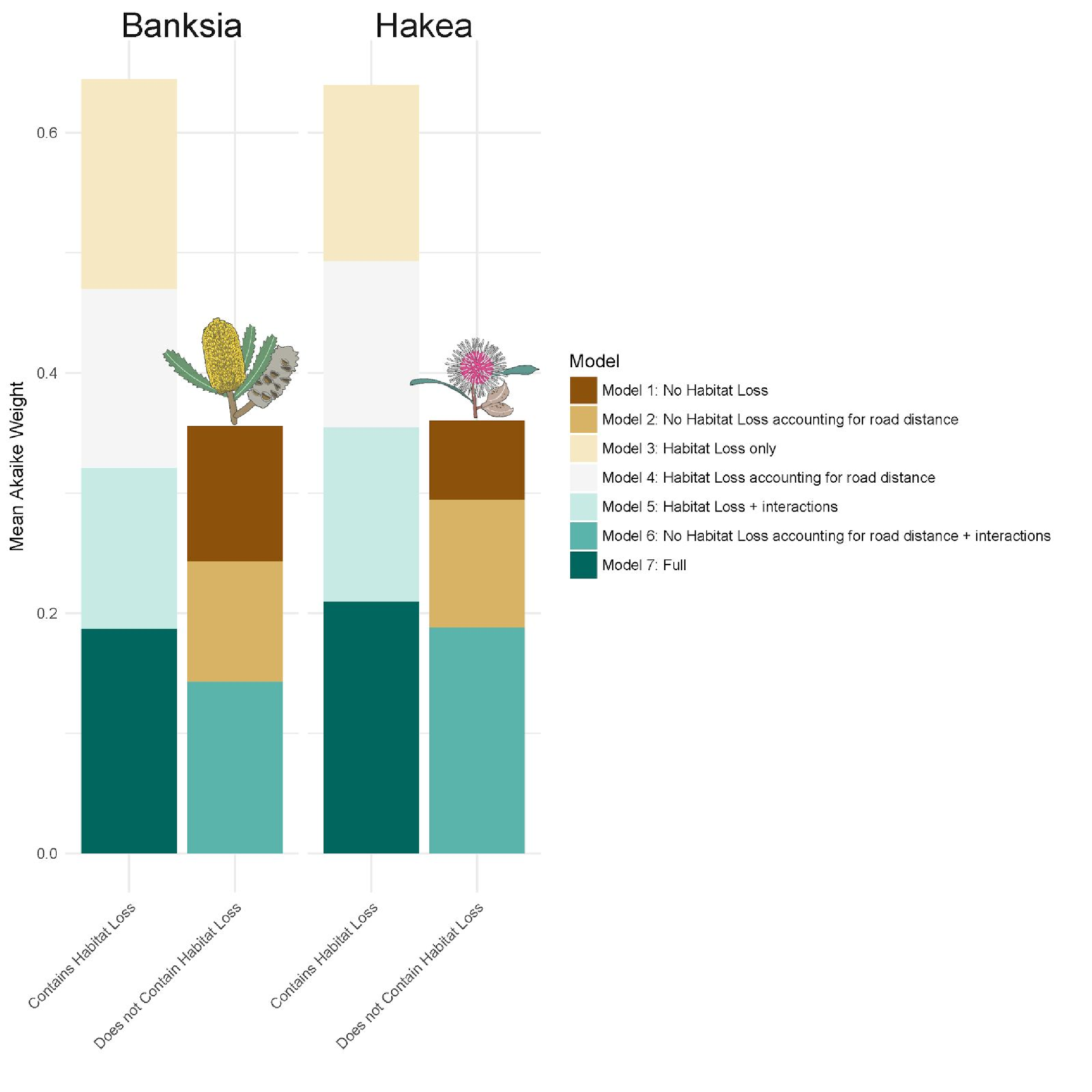
Summed Akaike weights for different PPM models run on *Banksia* and *Hakea* species. The height of each bar represents the mean Akaike weight for that type of model across all species. Higher Akaike weights correspond to better predictive models.

### Effect of Habitat Loss on Probability of Occurrence

We examined the effect of habitat loss by plotting the non-linear relationship between each variable and the density of occurrence implied by the coefficients estimated in the PPM for model 4 (*Habitat Loss accounting for Sampling Bias;* Figure 2), while setting all other coefficients to their mean (except the sampling bias variable which was always set to zero). For plotting we standardized both the mean probability of occurrence and habitat loss by dividing by their maximum values to make the relationships more easily comparable across species.To see how many species showed different classes of relationships with the bias proxy variables, we classified each relationship into positive, negative, or none (Figure 2). A relationship was classified as positive or negative if the correlation coefficient between mean probability of occurrence and a bias proxy variable was greater than 0.3 or less than −0.3 respectively (Figure 1, Figure 2).

**Figure 2.**
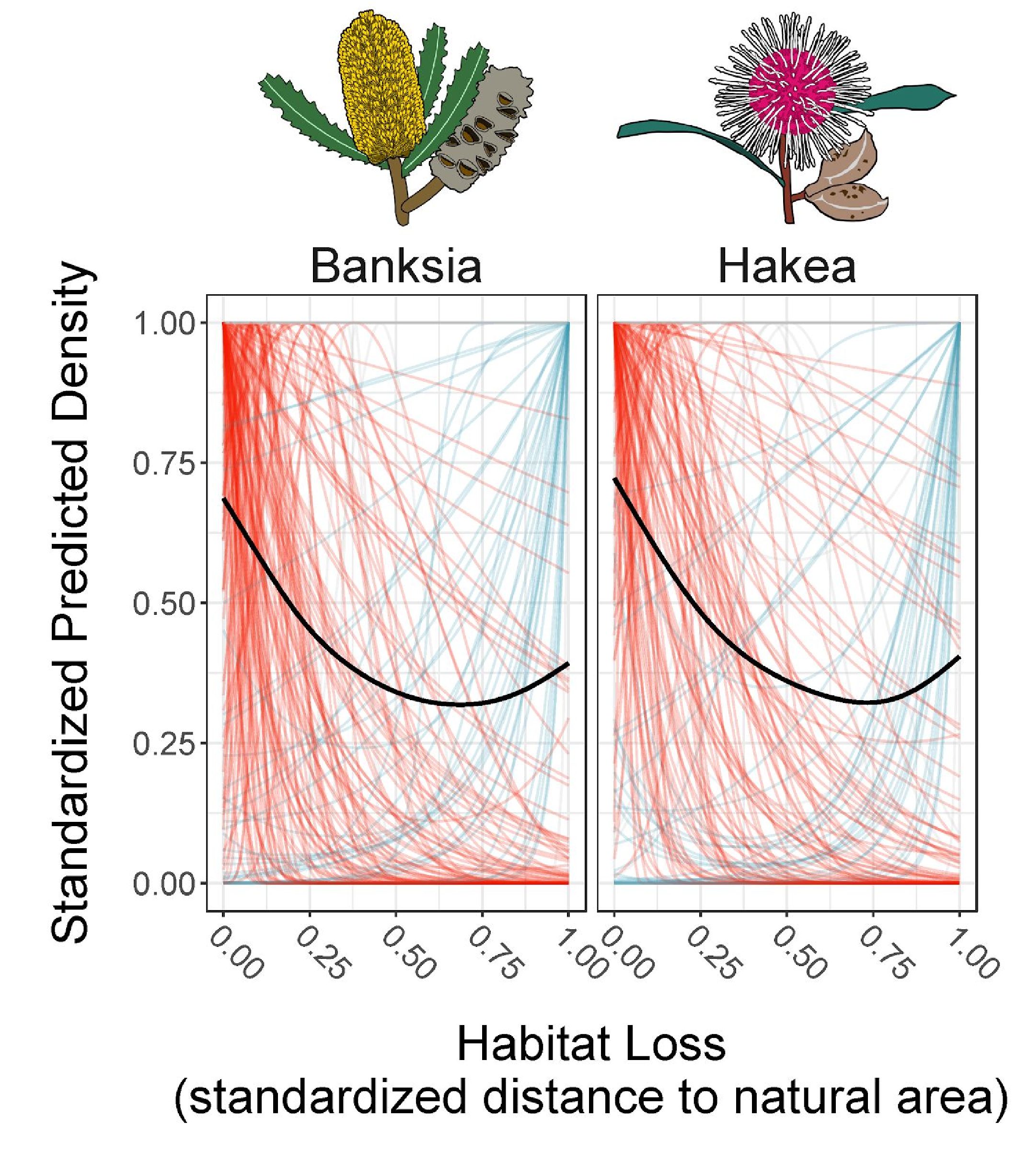
The relationships between distance to natural area (habitat loss proxy) and the density of occurrence for every modelled *Banksia* (left panels) and *Hakea* (right panels) species. Each line is the relationship for a different species. Lines are coloured red for a negative relationship, blue for a positive one, and grey for no relationship. The black line is a smoothed response showing the mean effect across all species shown in the panel. Both axes are standardized by dividing by the maximum value of each species, to make comparisons between species on a common scale.

### Effects of Incorporating Habitat Loss

We quantified the effects of incorporating habitat loss by comparing predictions made while accounting for habitat loss with those where it is unaccounted for. In order to control for habitat loss when predicting the ranges of *Banksia* and *Hakea* species, we can fix our bias proxy variables at a particular value (following (Warton et al. 2013)). Since we most likely observe a species if it actually occurs in areas with little or no habitat loss, it is reasonable to set the value of the habitat loss variable to zero when making predictions.We also set the sampling bias variable to zero (i.e. zero distance to a road), where sampling bias is least likely to occur. To explore the overall effects of fixing habitat loss during prediction, we predicted each species separately using the observed environmental variables and setting habitat loss to zero, or to its observed values. Each species was only predicted within those terrestrial ecoregions (Olson et al. 2001) from which at least one occurrence was recorded, to prevent biogeographically unrealistic predicted ranges that include suitable habitat far beyond the present-day extent of occurrence. For each species we used the best model that contained Habitat Loss, selected with AIC. All predictions were done at a resolution of 0.15° × 0.15°. Subsequently, we refer to the prediction made while not accounting for habitat loss as the “control predictions”, and those made while controlling for habitat loss as the “habitat loss predictions”.

### Effects of Incorporating Habitat Loss on Predicted Ranges

We quantified the effects of incorporating habitat loss on the overall size of predicted species ranges, as well as any shifts in their centre of density. We calculated the change in predicted range size by subtracting the total predicted density while accounting for habitat loss from the total predicted species density for the control predictions, and then standardising it by dividing by the mean of predicted total density for the control and habitat loss predictions.

We also calculated the centre of density of predicted species’ distributions by calculating the mean longitude and latitude of the predictions, weighted by the predicted density. The change in the centre of density was then visualized as a vector beginning at the control predictions and ending at the predictions that accounted for habitat loss (Figure 4).

### Effects of Incorporating Habitat Loss on Downstream Analyses

PPM models make predictions of occurrence point densities. In order to make inferences about species richness we need to convert these density estimates to predicted presences or absences. The PPM predicted density was conditioned on the total number of occurrence points, such that the total density of points predicted in the study area sums (approximately) to the total number of occurrence points (**δ**_tot_) in the dataset. To make different species’ predicted densities comparable, we standardized the predicted values by multiplying them by a species-specific constant. We inferred the presence of a species if the the poisson probability of observing at least one occurrence at the predicted density (1 – ɛ^−δ^) was greater than 0.05. If this probability was <0.05, we inferred an absence. To choose the multiplier, we calculated a value that would make the predicted total number of occurrence points for each species equal to the same value, that of the mean predicted number of occurrences across all species.

Next, we aggregated predicted numbers at a resolution of 0.15 degree square. We then divided Australia into 1.05 degree by 1.05 degree grids (7 × 0.15), and calculated the species diversity of *Banksia* and *Hakea* at this scale across Australia. We used multiplicative diversity partitioning to decompose the predicted diversity into Gamma, Alpha and Beta components. Gamma diversity corresponds to the total diversity within each 1.05 degree square grid cell. Alpha diversity is the mean diversity of each 0.15 degree square cell within the 1.05 degree square. Beta diversity is Gamma/Alpha. We used the R package entropart (Marcon and Hérault 2015) to calculate the diversity partitions.

We standardized all diversity values by subtracting the predicted diversity value under a model where all environmental variables and both bias variables (habitat loss and road proximity) were set to their observed values. This can be considered the control prediction, representing what we would predict without taking into account the difference between environmental and habitat loss variables.

We repeated the above analysis using the standardized predicted densities themselves in an abundance weighted version of the Alpha, Beta, and Gamma diversities, in order to see how accounting for habitat loss leads to shifts in the relative predicted densities across Australia.

We then tested the influence of accounting for habitat loss on the outcomes of Age-Range Correlations (ARC), a common macroevolutionary method based on the analysis of patterns of overlap in the inferred distributions of species. ARC quantifies the relationship between range overlap of species pairs and their age of divergence, to make inferences about the prevailing geographic mode of speciation within a clade (and other processes: see (Fitzpatrick and Turelli 2006, Warren et al. 2008)). We used the R package ‘phyloclim’ (Heibl and Calenge 2011) to calculate ARC for our control and habitat loss adjusted models for *Banksia* and *Hakea* separately, using phylogenies for *Banksia* from (Cardillo and Pratt 2013), and for *Hakea* from (Cardillo et al. 2017).

## Results

### Model Comparisons

Habitat loss was an important predictor of species recorded occurrences in most of the models. Summed Akaike weights show that models including the habitat loss proxy variables were on average nearly twice as likely to be the best model out of all the models tried (Figure 1). Out of the models that included habitat loss, all had nearly equal Akaike weight sums, suggesting that including interactions or road-related sampling bias often provided additional predictive power to the habitat loss models. (see Supplementary Figure 6 for a plot showing all interactions between habitat loss, sampling bias and environmental predictors in the full model). For subsequent analyses we compared the prediction from the best model including habitat loss, against the same model with habitat loss removed, on a species by species basis.

### Effect of Habitat Loss on Probability of Occurrence

The effect of habitat loss on the probability of species occurrence within our PPM models wasin the expected negative direction for most species. Most species of both genera experienced a steep decline in the probability of occurrence with increasing distance from natural areas (Figure 2). A small number of species however, showed the opposite trend, with strong increases in the mean probability of occurrence with increasing distance a natural area.

### Effects of Controlling for Habitat Loss on Species’ Predicted Ranges and Diversity

When we controlled for habitat loss by fixing its proxy variable at zero across the landscape, the predicted ranges of most *Banksia* and *Hakea* species were larger than when bias variables were not accounted for (Figure 3). Habitat loss had important effects on both genera, with large increases in predicted total density, both when sampling bias was additionally accounted for or not. This represents a mean increase of approximately 10% for species of *Banksia* and 12% for species of *Hakea*. For species occurring in the Southwest biodiversity hotspot (defined as any species with more than 50% of its predicted range occurring at a Latitude < −25 degrees, and a Longitude < 126 degrees), those numbers increased to 11% and 15% respectively.

**Figure 3.**
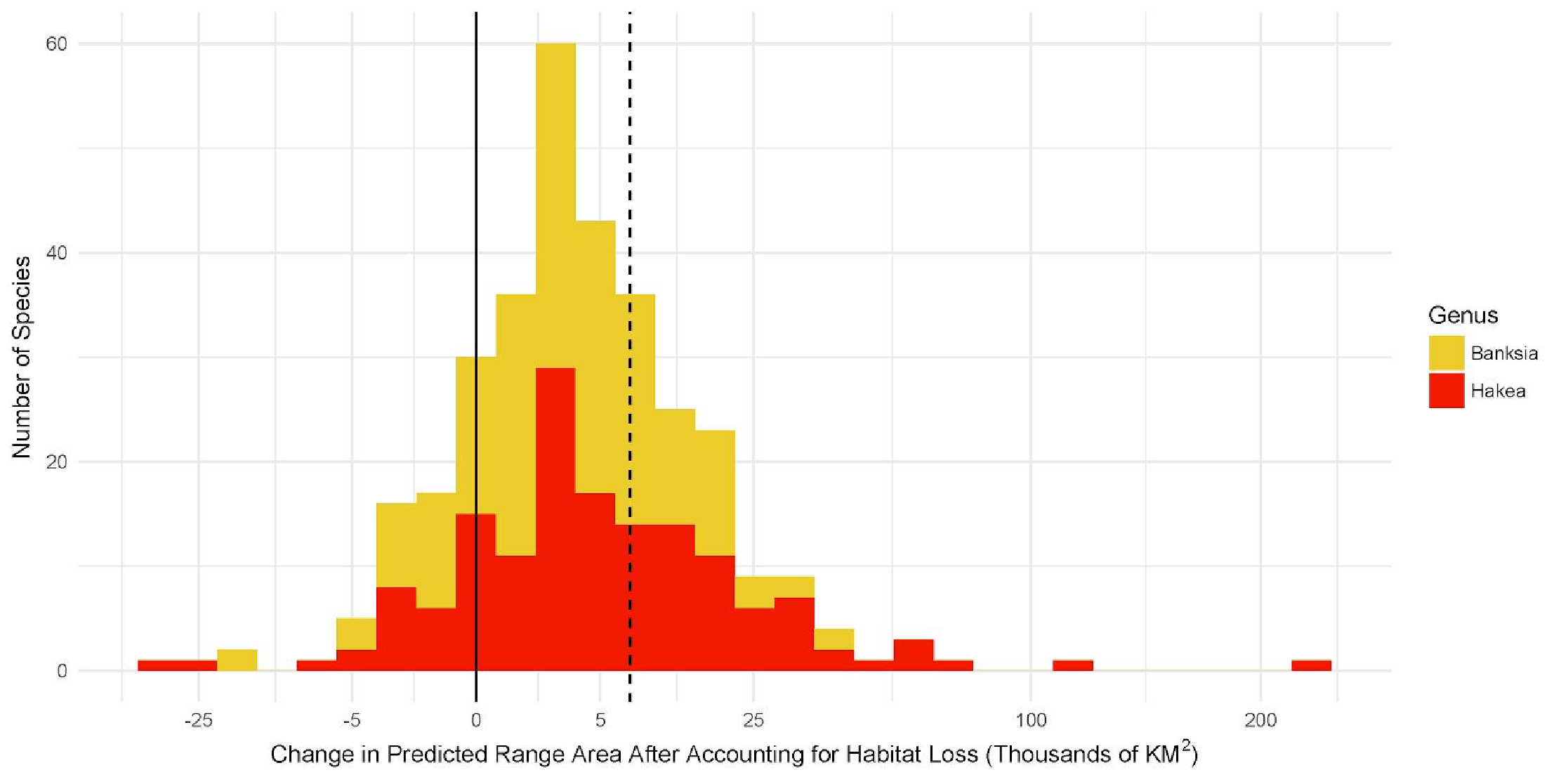
The change in the predicted range size between models accounting for habitat loss and those that do not. Each bar represents how many species fell in each interval on the × axis. The x axis represents the difference in predicted range size accounting for habitat loss, in thousands of kilometers squared, and split by genera; a positive value indicates that the predicted range size was larger when habitat loss was accounted for in the model. X axis is sign(x)sqrt(x) transformed. Vertical dotted line represents the mean difference ~ 7000 KM^2^).

For both *Banksia* and *Hakea*, gamma diversity and alpha diversity were higher when accounting for habitat loss, across most of the distribution of each genus (Figure 5). Beta diversity was mostly reduced by accounting for habitat loss. Using abundance-weighted diversity metrics showed similar patterns, although generally the results were more variable, with some regions that show mainly an increase in Alpha and Gamma diversity, now showing a decrease. We interpret these differences as a consequence of shifting density peaks for some species, leading to changes in evenness across the landscape. Indeed, when we plotted vectors showing how the centre of predicted density changed for *Banksia* and *Hakea* species, we found that for many species, accounting for habitat loss shifted the centre of density, sometimes substantially (Figure 4). It is clear from Figure 4 that the largest shifts occurred in the Wheatbelt region of the Southwestern Australian hotspot, where habitat loss has been particularly severe (Saunders 1989).

**Figure 4.**
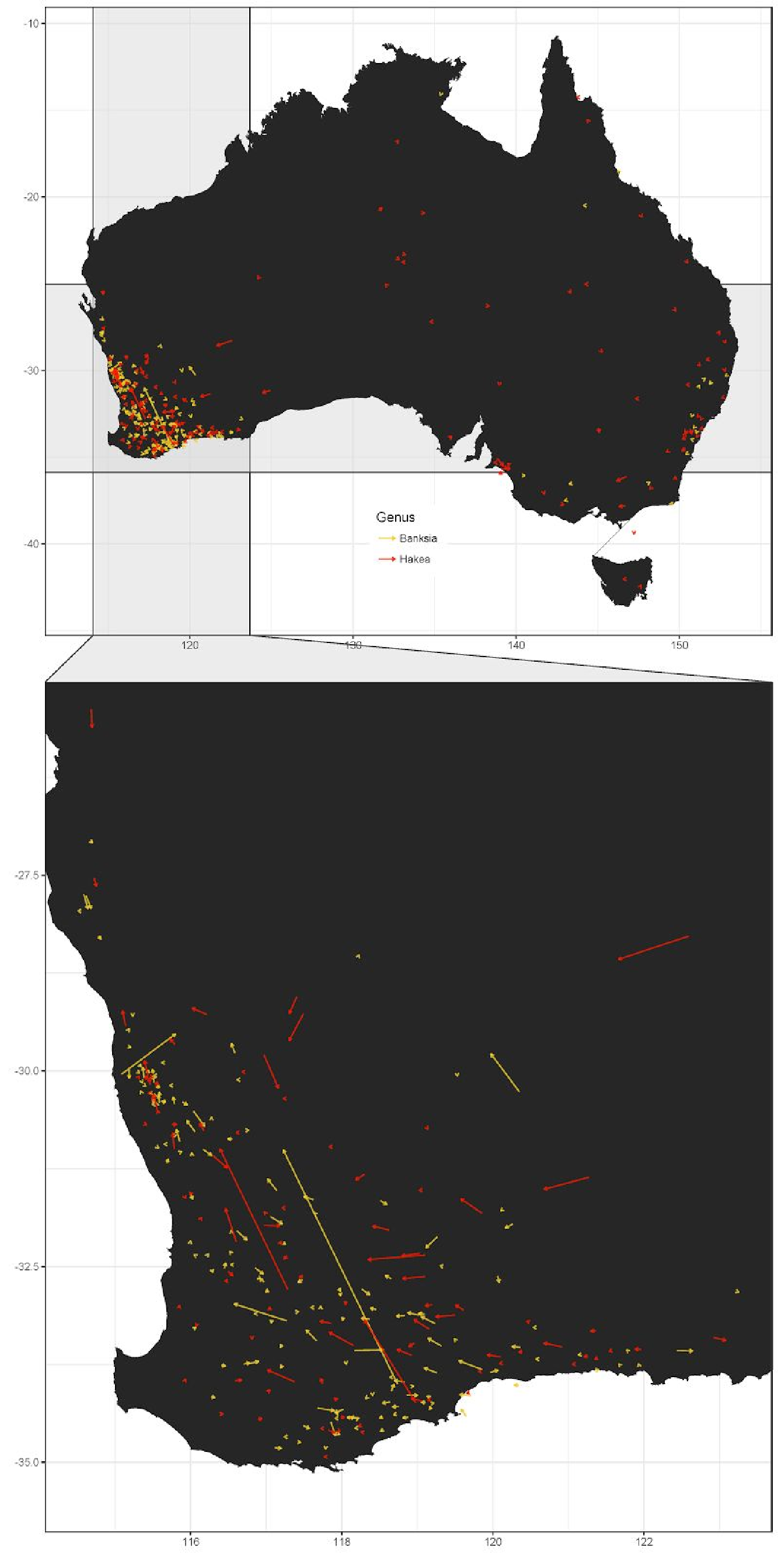
Map of changes in centre of predicted density for *Banksia* (red) and *Hakea* (yellow) species. The bottom of each arrow represents the predicted centre of density without accounting for habitat loss, the top (head) of each arrow is the predicted centre of density after accounting for habitat loss. Bottom panel is zoomed into the Western Australian biodiversity hotspot.

**Figure 5.**
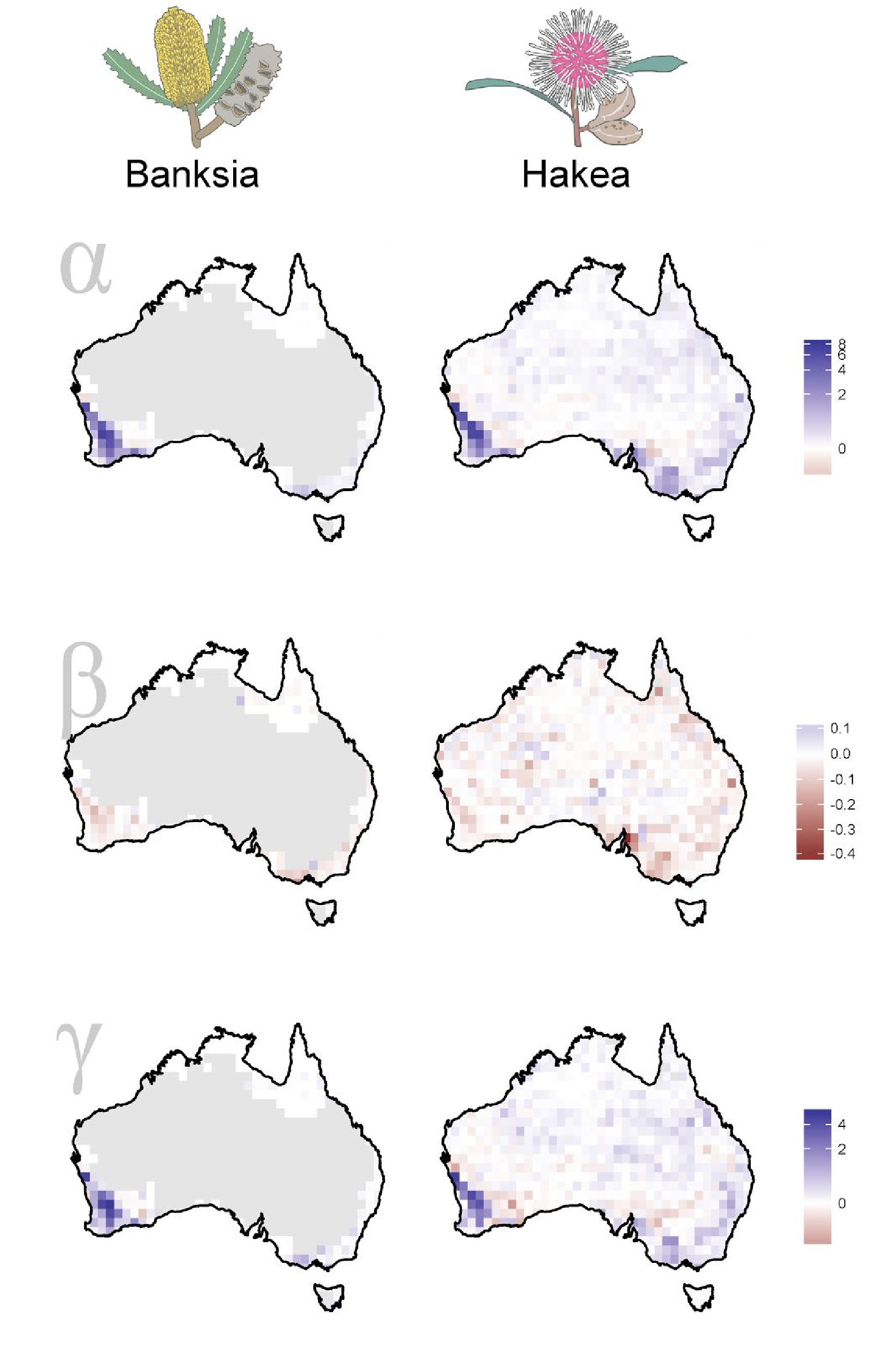
Maps showing the distribution of predicted diversity differences for *Banksia* (left column of panels) and *Hakea* (right column of panels) across Australia. Models controlling for habitat loss were used to predict individual species occurrences and the predictions were then summarised as diversity. Diversity was partitioned into Gamma (*γ*), Alpha (*α*), and Beta (*β*) diversity in 1.05 by 1.05 degree grid cells (based on sub-cells of 0.15 by 0.15 degree). Each diversity value was then standardized by subtracting the predicted diversity in a PPM model with the habitat loss proxy variable set to their observed values. Blue squares represent grid cells where diversity was higher when accounting for habitat loss, red squares where diversity values were lower. Grey squares were grid cells from regions in which there was not enough data to calculate diversity (e.g. less than 2 species). The colour scale is arcsinh transformed.

### Effects of Habitat Loss on Macroevolutionary Analyses

We conducted an Age-Range Correlation analysis (ARC), using predictions from the control and habitat loss models. Figure 7 shows that the slopes and intercepts of the associations between range overlap and divergence times of sister clades are not greatly changed by accounting for habitat loss, even though there are some large changes in predicted overlap among sister-clade pairs, mostly at recent divergence times. Most likely, the averaging procedure used to calculate overlap at deeper nodes cancels out much of the variation in overlap in opposite directions that we see at shallower nodes, so that the overall pattern is not strongly influenced by the changes in the predicted distributions of particular species.

## Discussion

Species Distribution Modelling was developed because species distributions inferred directly from occurrence records are likely to underestimate the real, potential, or historical extent of a species’ occurrence. Our results show that even modelled distributions are likely to underestimate species historical or potential distributions if they fail to account for the sampling bias introduced by historical habitat loss. Methods for inferring the potential distributions of species from their observed occurrences assume that occurrences are sampled without spatial bias: it is assumed that wherever a species occurs, it has been recorded. Our results confirm that failure to explicitly account for spatial biases in the distribution of occurrences reduces model performance: combined Akaike weights were only ~ 0.12 for *Banksia* and 0.08 for *Hakea* species, for models excluding both habitat loss and road distance. Our results show that the contraction of species distributions as a result of historical habitat loss introduces a bias with an influence on distribution models above and beyond that of the better-known sampling bias associated with road networks (Elith and Leathwick 2007, Phillips et al. 2009, Syfert et al. 2013, Warton et al. 2013, Fernández and Nakamura 2015, Fithian et al. 2015, Guillera-Arroita 2016). Furthermore, habitat loss modifies the predictive power of environmental variables for species distributions, sometimes in complex ways, since models including interactions with predictor variables tended to be better than models with main effects only. This implies that to more fully account for habitat loss or other biases, allowing interactions with other predictors and non-linear terms should be common practice, as suggested, for example, by Gelman (2007) for controlling for response bias in sociological survey analysis. For *Banksia* and *Hakea* species, distribution models that included habitat loss performed better, on average, than those that excluded habitat loss (Figure 1). Hence, most species distributions in these two genera, as inferred from recorded occurrences only, are spatially non-random subsets of the likely distributions prior to the recent, widespread conversion of much of southern and eastern Australia to anthropogenic landscapes. Given that *Banksia* and *Hakea* are two of Australia’s more well-known and conspicuous plant genera, the problem is likely to be at least as serious in other groups of plants.

On average, distribution models that accounted for habitat loss predicted distributions 10% and 12% larger than those that did not account for either sampling bias or habitat loss, for *Banksia* and *Hakea* respectively. For many species, however, this value was far higher: 22 species had predicted distributions >50% larger when habitat loss was accounted for. The obvious effect of larger predicted ranges on large-scale diversity patterns is to increase the number of overlapping species ranges within any given area - that is, to increase gamma and alpha diversity. When controlling for habitat loss, the models predict increases in gamma and alpha diversity across most of the regions in which *Banksia* and *Hakea* are distributed (Figure 5). Of more concern, though, is that the influence of habitat loss on predicted diversity is geographically heterogeneous. Unsurprisingly, the influence of habitat loss on gamma and alpha diversity is minimal across inland and northern Australia, where habitat loss has been limited. The effect of habitat loss on both diversity (Figure 6) and mean predicted range size is most profound in the Southwest Australian biodiversity hotspot. This probably results from both of the criteria that qualify this region as a global biodiversity hotspot: the extensive conversion of natural habitat to agricultural landscapes, and the high proportion of narrowly-endemic species known only from very restricted areas. Spatial turnover in species composition (beta diversity) shows patterns that are largely the converse of alpha and gamma diversity: there is a general pattern of decreased beta diversity. Because alpha and beta diversity were calculated by multiplicative partitioning of gamma diversity, this suggests that on average, the increases in alpha diversity under bias-explicit models were greater than the increases in gamma diversity.

**Figure 6.**
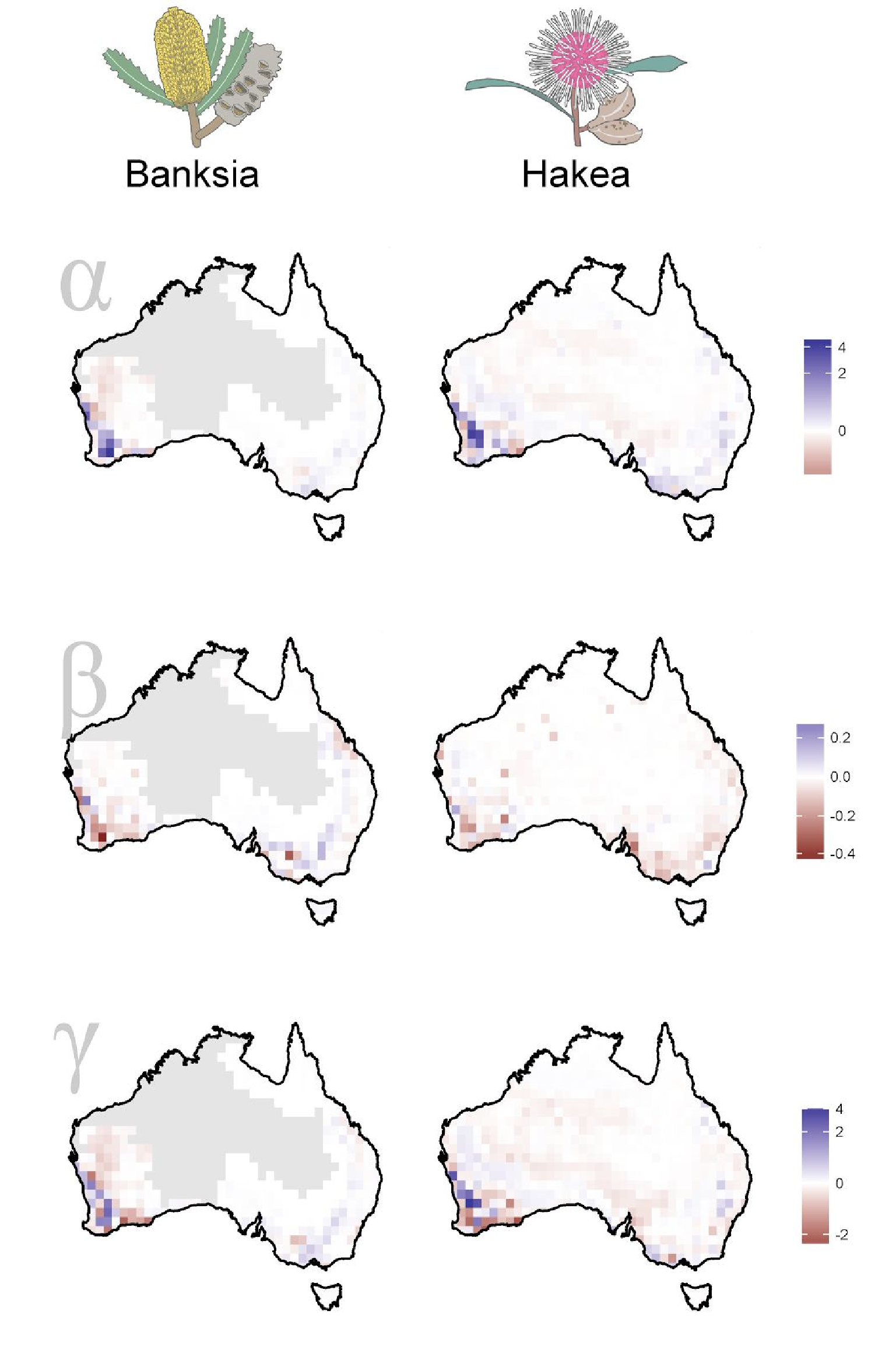
Maps showing the distribution of abundance-weighted predicted diversity differences for *Banksia* (left column of panels) and *Hakea* (right column of panels) across Australia. Models controlling for habitat loss were used to predict individual species densities and the predictions were then summarised as abundance-weighted diversity (where predicted density was treated at abundance). Diversity was partitioned into Gamma (γ), Alpha (α), and Beta (β) diversity in 1.05 by 1.05 degree grid cells (based on sub-cells of 0.15 by 0.15 degree). Each diversity value was then standardized by subtracting the predicted diversity in a PPM model with the habitat loss proxy variable set to their observed values. Blue squares represent grid cells where diversity was higher when accounting for habitat loss, red squares where diversity values were lower. Grey squares were grid cells from regions in which there was not enough data to calculate diversity (e.g. less than 2 species). The colour scale is arcsinh transformed.

**Figure 7.**
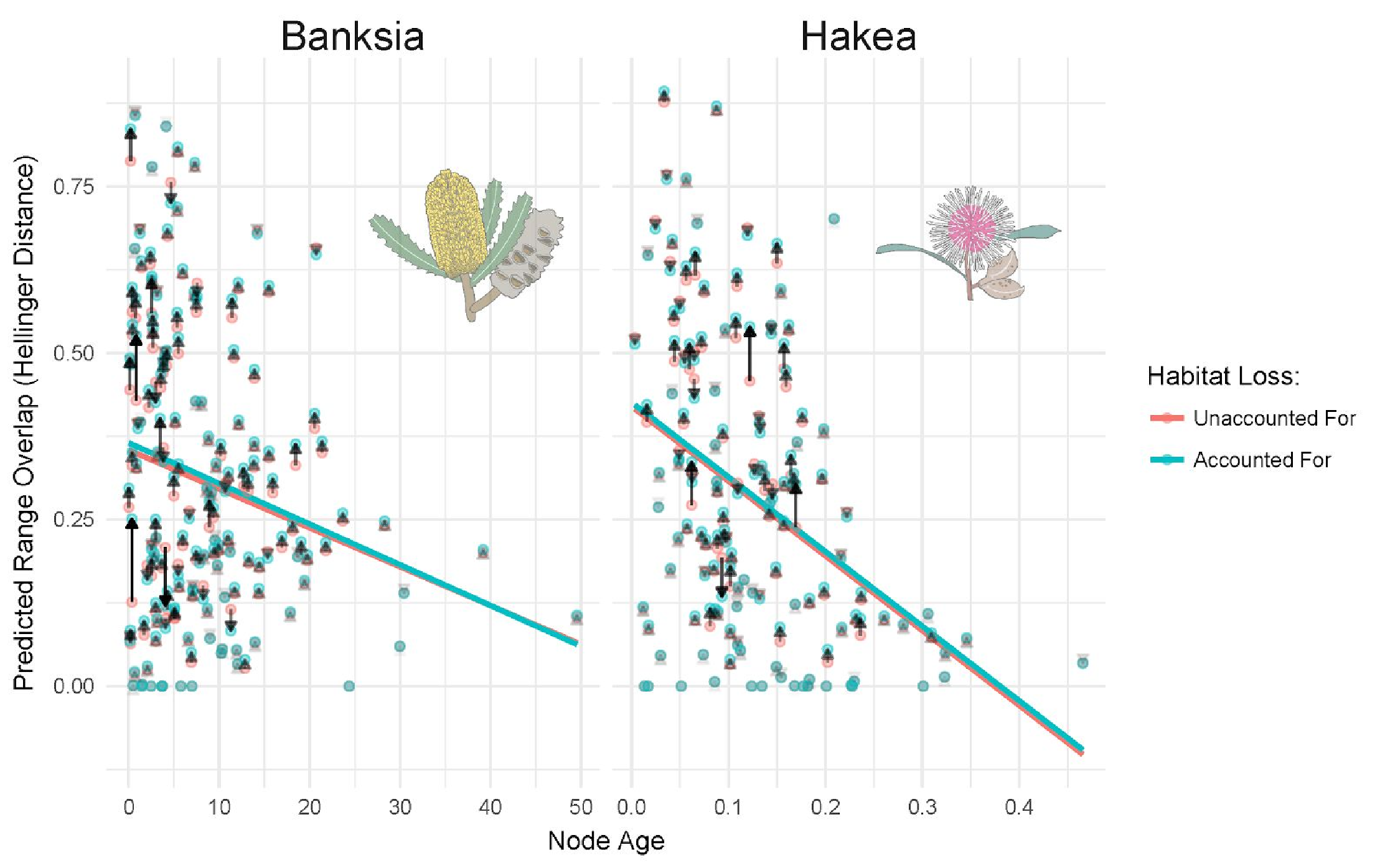
Age of nodes versus predicted range overlap for *Banksia* and *Hakea* species. Range overlap was calculated using predicted densities from either a control model (red), or a model accounting for habitat loss (blue). Arrows are drawn from the control predictions to the habitat loss accounted predictions, with transparency proportional to the overall change. Lines are lines of best fit under a simple linear model.

Our approach to incorporating habitat loss into the predictive models was to set the value of its proxy variables to zero, as this is the value for which we expect the least reduction in observation probabilities. However, this requires us to assume that the habitat loss proxy variables entirely reflect bias in the sampling of species occurrences, and do not have any genuine, biological influence of their own on species occurrences. In general, this is probably a reasonable assumption, but there may be some cases in which this assumption is violated. For example, the distribution of roads across a landscape may not always be random with respect to environmental features: in some regions, roads are more likely to traverse slightly elevated areas with well-drained soils than low-lying, flood-prone areas, and to follow moderate contours across the landscape rather than steep slopes. Each of these environments may have different soil types that differ in nutrient status, pH or moisture holding capacity. Species that either favour or cannot tolerate such soils will have distributions of occurrence records that are positively or negatively associated with roads, for genuine biological (albeit indirect) reasons. In the same way, the degree of historical habitat loss is also not random with respect to soil type and other features of the environment. In regions of southern and southwestern Australia now largely under wheat cropping, areas of sandy soils with low moisture-holding capacity are more likely to retain natural habitat than areas with soils more favourable for cropping. Although such cases of genuine association between species occurrence and bias proxies are likely to be uncommon, they make it difficult to be certain that our proxies are fully accounting for bias and not weakening or reinforcing genuine patterns of occurrence. One way to tease apart the bias and biological components of a variable‘s association with species occurrences may be to use models that combine presence-only data with presence-absence data (Fithian et al 2015), although we do not yet have systematically-collected presence-absence data that includes all species of *Banksia* and *Hakea*. However, it is almost certainly preferable to estimate and account for biases imperfectly than to infer species distributions without considering biases at all.

Our results also suggest that a further reason to explicitly account for biases in distribution models is that the biases interact with environmental variables in the models. This means that not only are distributions inferred directly from occurrences an underestimate (in most cases) of modelled distributions, but the climatic and habitat types within the directly inferred distributions may differ from those within the modelled distributions. The complex nature of the interactions means that it is difficult to generalize about this kind of qualitative difference between directly inferred and modelled ranges. Nevertheless, this could have significant effects on some downstream ecological and evolutionary analyses of species environmental niches or responses to future environmental change. In sociology, it is well-known that variables related to biases in, for example, survey response, should be incorporated into regression models, and that interactions with predictors should also be included when possible (Gelman 2007).

Though we compared the effect of habitat loss to that of the effect of sampling bias on observations of species, they do differ in a number of important ways. Sampling bias affects the probability of a species that is currently in a region being observed. For this reason, it is likely best practice to try and account for it in all cases. On the other hand, habitat loss primarily imposes a bias in the probability of observing a species that historically existed in an area, but is genuinely no longer present. This means that whether or not habitat loss should be accounted for in SDMs will depend on the goals of the study. If the goal is to best predict where a species is currently found, then habitat loss should be included in the model, with its observed values used during prediction (rather than setting the values to zero). This is because for many species habitat loss will be highly informative in predicting where a species is not currently found. On the other hand, for many other kinds of questions it is of more relevance to understand the extent of species historical distributions, as the basis for understanding the non-anthropogenic ecological, evolutionary and biogeographic processes that shape species distributions.

Inaccurate estimates of species distributions, and by extension, of spatial patterns of diversity, may limit or bias our inference of ecological or evolutionary processes. Age-Range Correlation, for example, is based upon analysis of the overlap in species’ distributions with respect to their divergence times (e.g. (Phillimore et al. 2008, Schnitzler et al. 2011, Anacker and Strauss 2014, Cardillo and Warren 2016). In our analysis, we found that SDM models accounting for habitat loss increased the estimated degree of overlap among species, but did not appreciably alter the intercept or slope of the ARC. As we pointed out earlier, however, this is likely to be due to the “averaging-out” of the overlap patterns at deeper nodes in the phylogeny, under the ARC method we used (Fitzpatrick & Turelli 2006), which implies that there was little phylogenetic signal in the impact of habitat loss on predicted species’ range for *Banksia* and *Hakea*. Other approaches to reconstructing geographic modes of speciation, such as the proportion of sympatric sister species (e.g. (Phillimore et al. 2008, Schnitzler et al. 2011, Anacker and Strauss 2014, Cardillo and Warren 2016), Cardillo & Warren 2016), are based only on shallow phylogenetic divergences and may be more sensitive to changes in overlap patterns that result from accounting for habitat loss in SDMs.

A loss of accuracy in predicting species ranges due to habitat loss biases could also be a problem for species and ecosystem conservation. Species range estimates are often used in making decisions about species conservation, such as in the design of reserves, climate change planning, and invasive species modelling (Rodríguez et al. 2007, Guisan et al. 2013). The issues of habitat loss biases could be particularly problematic for conservation because those species that have experienced the highest amount of habitat loss, and are therefore likely to be in the most need of conservation efforts, are also the most likely to have the accuracy of their range predictions affected. Therefore, being able to at least estimate, and to some extent control for the effect of habitat loss on species range reconstructions, as we have done here, could be very important for improving the use of species distribution models in conservation planning.

## Data Availability

Data used in this study can be retrieved from the Atlas of Living Australia. Occurrence records were cleaned prior to analysis by removing obviously incorrect records (such as those who coordinates referred to the herbarium where the sample was housed, rather than where it was collected from). Cleaned data is available upon request from the authors.

## Acknowledgements

We’d like to thank Anna Simonsen for thoughtful comments on a previous version of the manuscript, and Ian Renner for helpful advice and conversation on using PPM models. This work was funded by Australian Research Council Discovery Grant DP110103168.

## Supplement

### Supplementary Methods 1: Calculating a Continuous Habitat Loss Measure

To quantify habitat loss we downloaded raster data on dynamic vegetation land cover from Geoscience Australia (http://www.ga.gov.au/scientific-topics/earth-obs/landcover), and converted it to a binary variable where each grid cell in Australia was classified as ‘natural’ (habitat intact) or ‘unnatural’ (habitat lost) vegetation. This dataset classified each 250m square grid cell in Australia into 22 vegetation classes, using satellite reflectance data to train a predictive machine learning model (Dynamic Markov Chain modelling). There were 12 categories we considered natural, and 8 categories we considered unnatural (see Supplementary Table 1). Bodies of water and salt lakes were excluded from the analysis.

One potential problem with categorical data in general, and binary data particularly, is that imprecision or inaccuracy in spatial coordinates of occurrence records can result in the opposite vegetation state becoming associated with a record, if it is incorrectly recorded as being outside the boundary of the correct grid square. In order to reduce this problem, and to make this variable more comparable to our sampling bias proxy – which was the geographic distance to the nearest road (Supplementary Figures 1 and 2) – we created a proximity map based on area classified as ‘natural‘. Using QGIS (Quantum GIS Development, 2015) we created a raster that calculated the distance in each 100X100m square of Australia to the nearest area classified as “natural”. Therefore, points falling within a natural square were assigned a value of 0, with values increasing with greater distance from a natural square. Supplementary Figures 3 and 4 show maps of this variable across Australia, and Southwestern Australia respectively.

### Supplementary Methods 2: Point Process Models, and Variable Selection

#### Point Process Models

To model the individual distributions of each species, we used point-process models (PPM; (Renner *et al.*, 2015). PPMs model occurrence points as a spatial Poisson process, where the density of occurrences at any given spatial point is distributed according to a continuous Poisson distribution, whose intensity value is a linear function of the environmental variables. PPMs can accommodate both linear and polynomial terms, and interactions between variables. PPMs have been shown to be a generalization of several other well-known modelling frameworks, including general linear models and MAXENT (Renner & Warton, 2013). At the core, PPMs are linear models, so in order to model more complicated relationships between environment and occurrence, polynomial combinations of variables are created. Among other consequences, this means that the total number of variables to include in the model is limited, because the number of polynomial combinations increases exponentially with the number of variables. In order to keep the number of variables in our models reasonable, we first used a variable selection step to choose the top 8 explanatory variables for each species.

#### Model Selection

We began with a set of 104 environmental variables from the Atlas of Living Australia (Supplementary Table 2). To select environmental variables from this set that were highly associated with each species’ distribution we ran separate correlation clustering and Guided Regularized Random Forest (GRRF) analyses on each species (Deng & Runger, 2013).

##### Correlation Clustering

We removed highly correlated variables for each species, by conducting hierarchical clustering on all 104 variables based on their Pierson‘s correlation, defining clusters by cutting the clustering tree at a correlation of 0.75, and then only keeping one variable from each cluster – the variable in the cluster with the highest importance score calculated in the next step, using GRRF.

##### Guided Regularized Random Forest (GRRF)

This method uses random forest for classification, a machine learning method that uses ensembles of classification trees (or regression trees for continuous responses) for prediction (Breiman, 2001). In this case we are classifying points from our full dataset as either occurrence points for a particular species, or as ‘background’ points, using the same quadrature points we use for the PPM as background points, using all possible environmental variables as predictors. GRRF works by first running a regular random forest on the data. From this first random forest, importance scores are generated for each explanatory variable, reflecting how much each one contributes overall to the *out of bag* (OOB) estimates (based on how well the model predicts held back samples). A second random forest is then run on the same data, but this time variables are weighted by their importance when deciding on the best splits. This has the effect of regularizing the variables, so that the algorithm preferentially chooses the better overall predictor in cases where two or more variables have similar explanatory power at a particular split. After filtering the variables by choosing only the variables with the highest GRRF importance scores within each correlation cluster (see Correlation Clustering), we then further filtered the variables, by choosing only the environmental variables with the highest eight importance scores, excluding our bias variables. In the PPM analysis, we used these best eight variables plus our two bias variables for a total of 10 variables for each species. To highlight the influence of the two bias variables, we also constructed PPMs using only the eight environmental variables, and compared these with the 10-variable models.

---

**Supplementary Table 1.** Classes from the dynamic vegetation land cover dataset from Geoscience Australia. Each class was determined to be natural or not natural, as specified in the second column, for further analysis.

| **Class** | **Natural?** |
| --- | --- |
| **Closed Tussock Grassland** | Yes |
| **Open Tussock Grassland** | Yes |
| **Open Hummock Grassland** | Yes |
| **Scattered shrubs and grasses** | Yes |
| **Dense Shrubland** | Yes |
| **Open Shrubland** | Yes |
| **Closed Forest** | Yes |
| **Open Forest** | Yes |
| **Woodland** | Yes |
| **Open Woodland** | Yes |
| **Rain fed cropping** | No |
| **Irrigated cropping** | No |
| **Rain fed sugar** | No |
| **Irrigated sugar** | No |
| **Rain fed pasture** | No |
| **Irrigated pasture** | No |
| **Mines and Quarries** | No |
| **Cities and towns** | No |
| **Wetlands** | Yes |
| **Alpine meadow** | Yes |
| **Lakes and dams** | Excluded |
| **Salt lakes** | Excluded |

**Supplemental Table 2.** Atlas of Living Australia id and description for 104 variables used in SDM models for *Banksia* and *Hakea*. Some were later discarded due to too much missing data. Variables with “(BioXX)” are from the bioclim dataset.

| id | description |
| --- | --- |
| cl1053 | Terrestrial Ecoregional Boundaries |
| el1077 | Gross Primary Productivity (2012-03-13) |
| el1078 | Normalized Difference Vegetation Index (2012-03-05) |
| el1079 | Enhanced Vegetation Index (2012-03-05) |
| el1080 | Fraction of Photosynthetically Active Radiation (fPAR) |
| el1081 | Leaf Area Index (LAI) - 2012-03-05 |
| el2016 | Euclidean Distance to Coast (metres) |
| el597 | Erosivity |
| el603 | Nitrogen |
| el607 | Dolomite (physical) |
| el608 | Gypsum |
| el609 | Potassium |
| el610 | Dolomite (acidity) |
| el615 | Phosphorus |
| el616 | Lime |
| el661 | Erodibility |
| el662 | Water holding capacity |
| el663 | ph |
| el665 | Carbon - organic |
| el666 | Drainage - variability |
| el667 | Moisture - variability |
| el668 | Evaporation - average |
| el669 | Moisture - average |
| **el670** | Drainage - average |
| **el671** | Evaporation - variability |
| **el672** | Runoff - average |
| **el673** | NDVI Mean |
| **el674** | Elevation |
| **el675** | Slope length |
| **el676** | NPP Mean |
| **el680** | Aspect |
| **el728** | Humidity - annual mean relative |
| **el806** | Bulk density |
| **el807** | Lithology - mean age (log) |
| **el808** | Lithology - mean age |
| **el809** | Lithology - mean age range (log) |
| **el810** | Distance - to any water (weighted) |
| **el811** | Phosphorus - plant-available pre-European |
| **el812** | Water holding capacity - plant-available |
| **el814** | Clay % |
| **el815** | Valley bottom % |
| **el816** | Soil depth |
| **el817** | Phosphorus pre-European |
| **el819** | Distance - to coast |
| **el820** | Distance - to non-permanent water (weighted) |
| **el821** | Nitrogen concentration pre-European |
| **el822** | Nitrogen store pre-European |
| **el824** | Magnetic anomalies |
| **el825** | Pedality hydrological score |
| **el826** | Topographic wetness index |
| **el827** | Topographic relief |
| **el828** | Phosphorus - plant-available total pre-European |
| **el829** | Calcrete |
| **el830** | Distance - to permanent water (weighted) |
| **el831** | Nitrogen - plant-available pre-European |
| **el832** | Bouguer gravity anomalies |
| **el833** | Hydrologic conductivity - average saturated |
| **el834** | Ridge top flatness |
| **el835** | Lithology - age range |
| **el836** | Topographic slope (%) |
| **el837** | Carbon store pre-European |
| **el838** | Lithology - fertility |
| **el840** | Valley bottom flatness index |
| **el841** | Soil pedality |
| **el844** | Topographic roughness |
| **el845** | Soils - coarse |
| **el849** | Wind speed - annual mean 9am |
| **el859** | Wind speed - annual mean 3pm |
| **el860** | Wind run - annual mean |
| **el861** | Radiation - driest quarter (Bio25) |
| **el862** | Temperature - annual range (Bio07) |
| **el863** | Precipitation - coldest quarter (Bio19) |
| **el864** | Moisture Index - lowest quarter mean (Bio33) |
| **el865** | Moisture Index - highest quarter mean (Bio32) |
| **el866** | Precipitation - wettest period (Bio13) |
| **el867** | Temperature - coldest period min (Bio06) |
| **el868** | Moisture Index - warmest quarter mean (Bio34) |
| **el869** | Radiation - coldest quarter (Bio27) |
| **el870** | Temperature - wettest quarter mean |
| **el871** | Radiation - lowest period (Bio22) |
| **el872** | Precipitation - driest period (Bio14) |
| **el873** | Moisture Index - coldest quarter mean (Bio35) |
| **el874** | Temperature - annual mean (Bio01) |
| **el875** | Temperature - driest quarter mean (Bio09) |
| **el876** | Temperature - coldest quarter mean (Bio11) |
| **el877** | Radiation - wettest quarter (Bio24) |
| **el878** | Precipitation - warmest quarter (Bio18) |
| **el879** | Temperature - warmest period max (Bio05) |
| **el880** | Radiation - highest period (Bio21) |
| **el881** | Radiation - annual mean (Bio20) |
| **el882** | Precipitation - seasonality (Bio15) |
| **el883** | Temperature - isothermality (Bio03) |
| **el884** | Moisture Index - highest period (Bio29) |
| **el885** | Moisture Index - seasonality (Bio31) |
| **el886** | Precipitation - wettest quarter (Bio16) |
| **el887** | Radiation - seasonality (Bio23) |
| **el888** | Temperature - diurnal range mean (Bio02) |
| **el889** | Precipitation - driest quarter (Bio17) |
| **el890** | Temperature - warmest quarter (Bio10) |
| **el891** | Moisture Index - annual mean (Bio28) |
| **el892** | Temperature - seasonality (Bio04) |
| **el893** | Precipitation - annual (Bio12) |
| **el894** | Radiation - warmest quarter (Bio26) |
| **el895** | Moisture Index - lowest period (Bio30) |

**Supplementary Figure 1.**
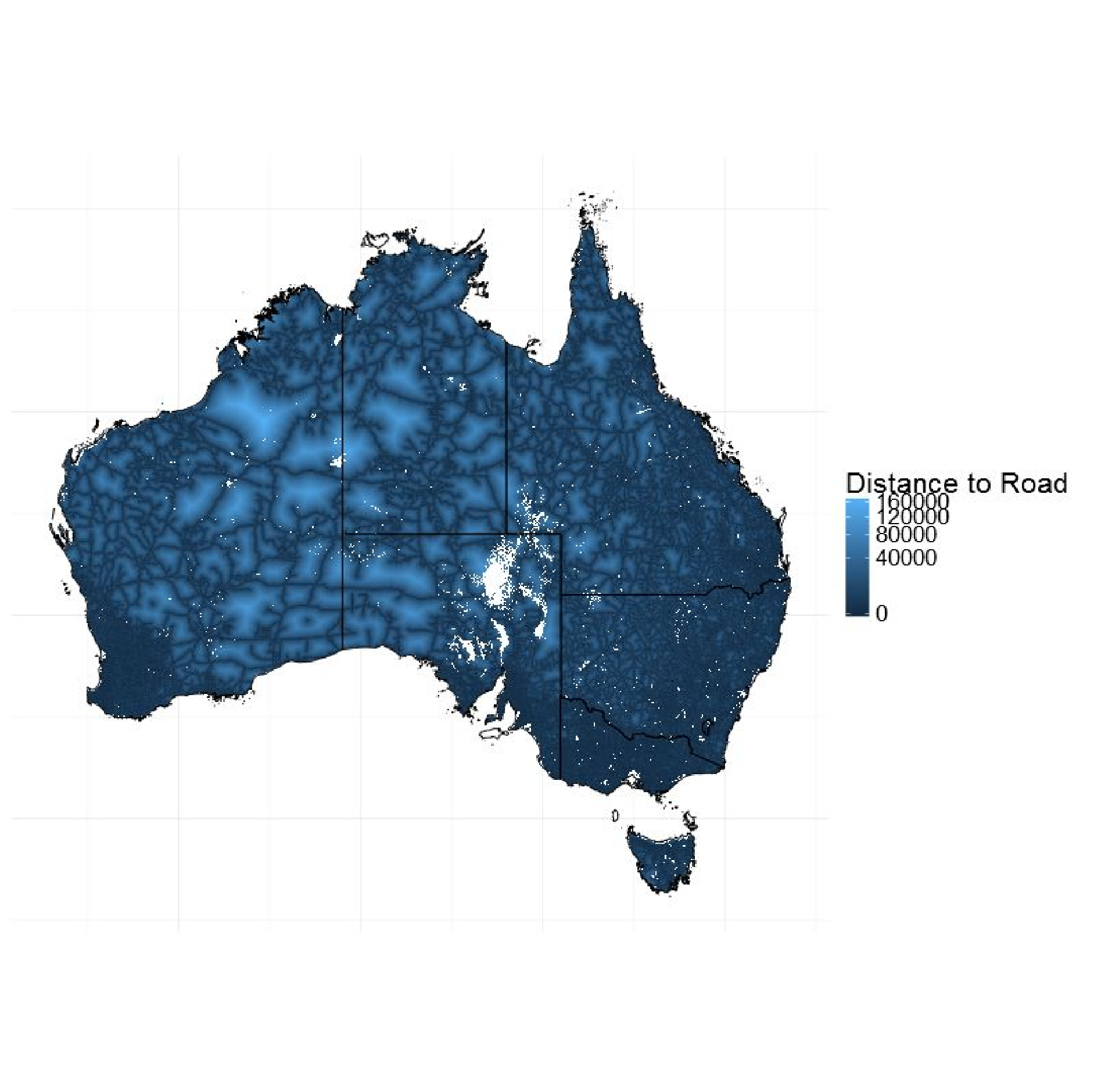
Map of distance to road sampling bias proxy variables across Australia. Colour bar is square root transformed.

**Supplementary Figure 2.**
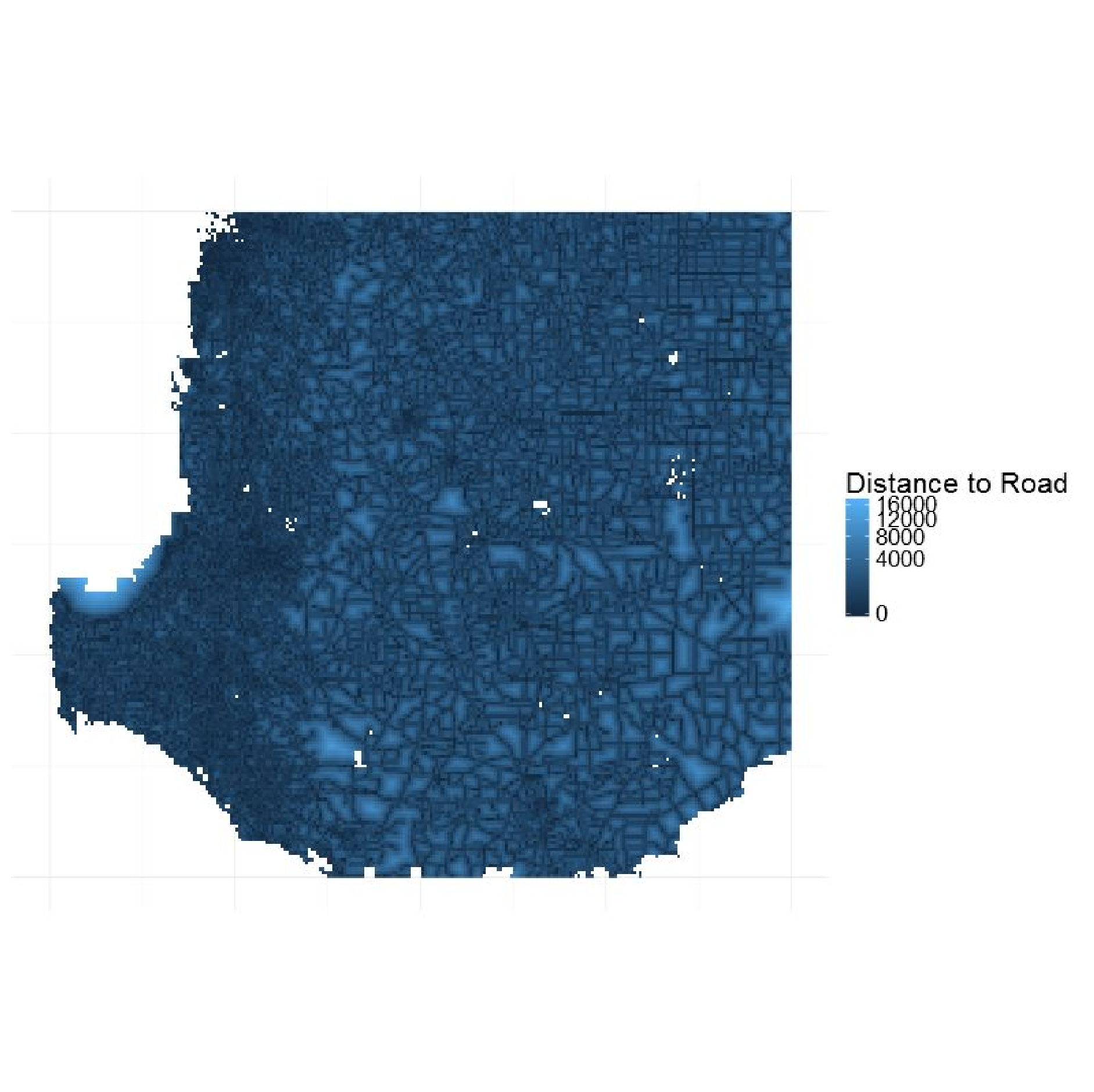
Map of distance to road sampling bias proxy variables for Southwestern corner of Australia, where Proteaceae reaches its diversity peak. Colour bar is square root transformed.

**Supplementary Figure 3.**
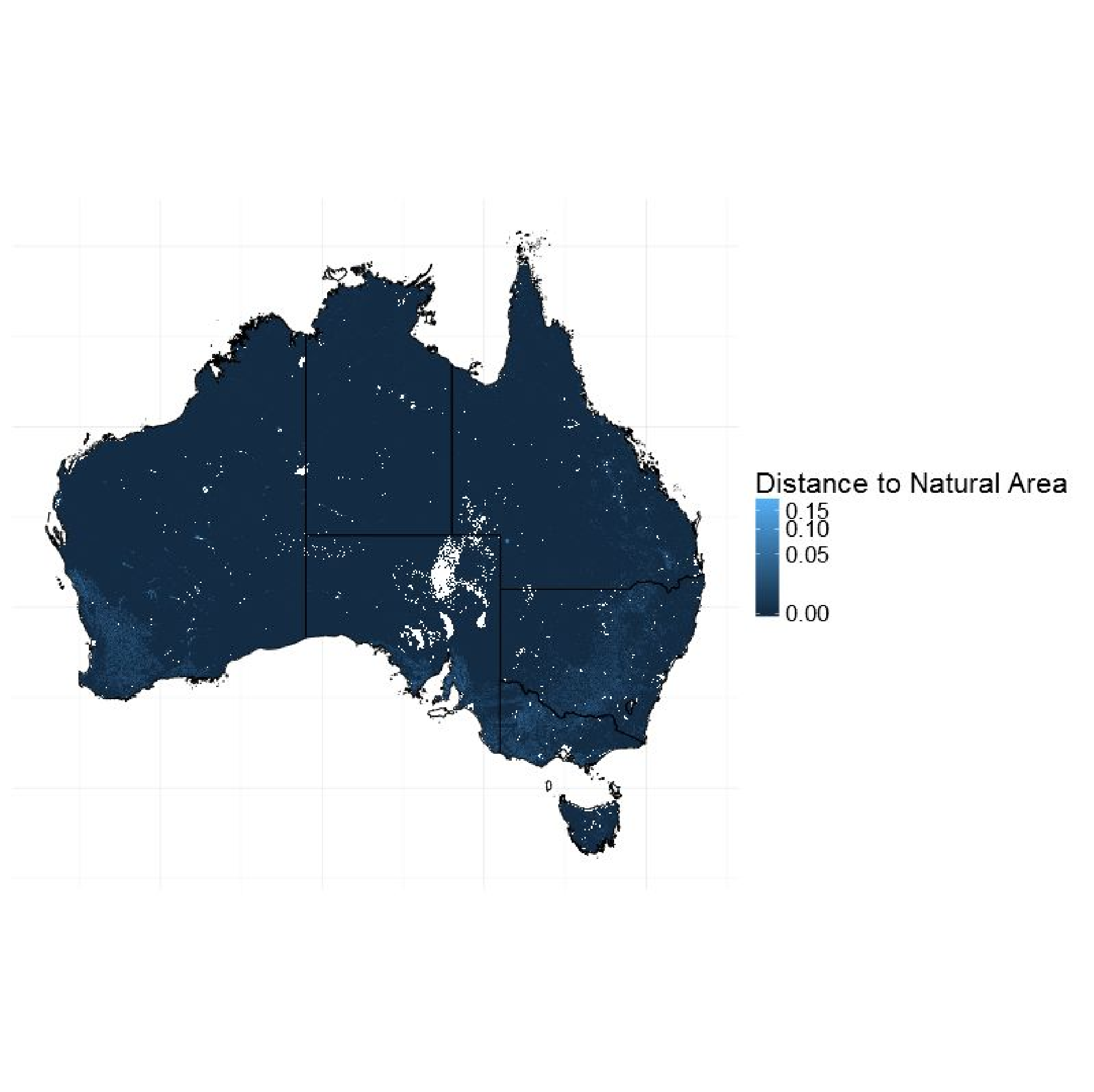
Map of distance to natural area habitat loss bias proxy variable across Australia. Colour bar is square root transformed.

**Supplementary Figure 4.**
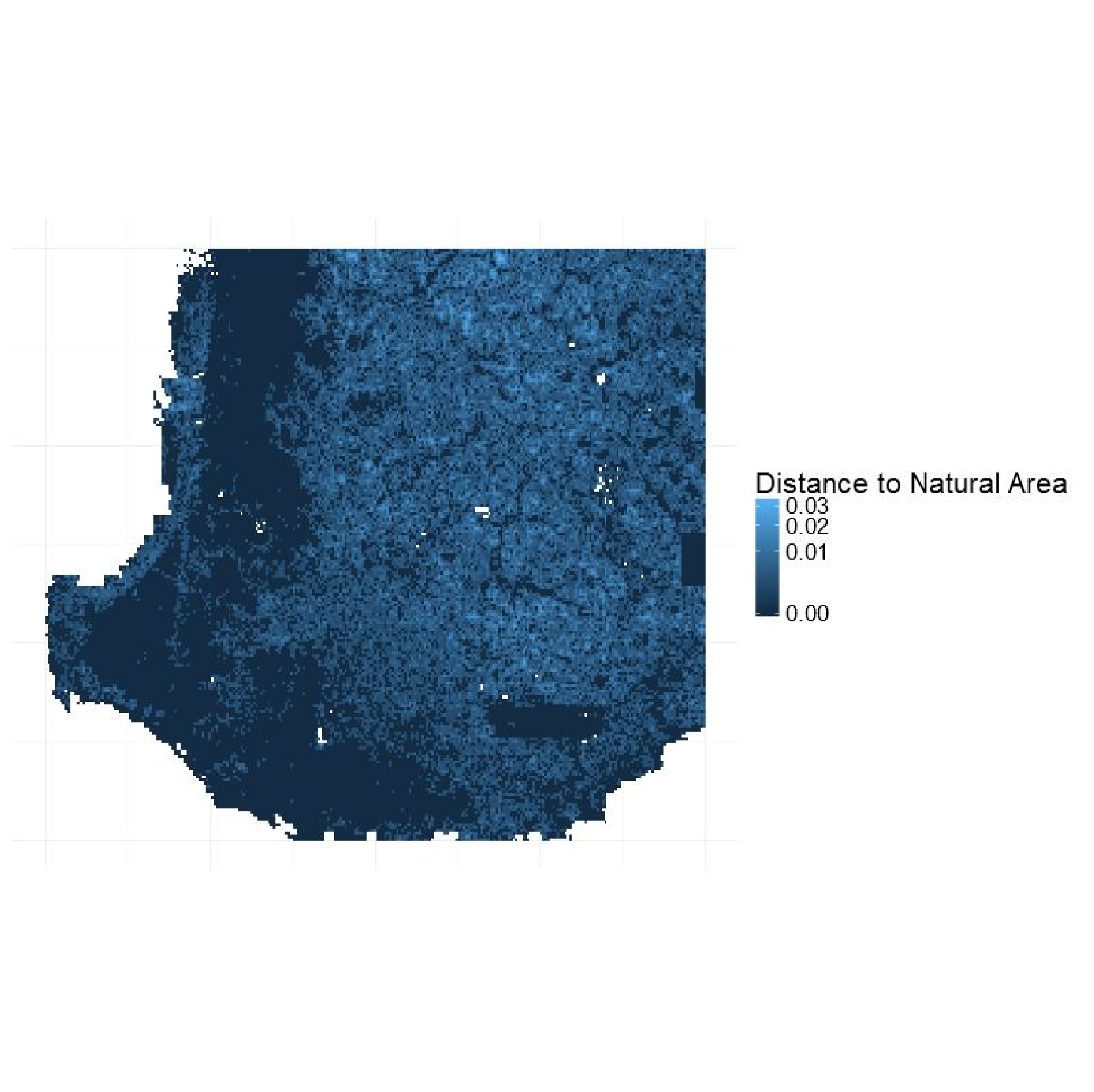
Map of distance to natural area habitat loss bias proxy variables for Southwestern corner of Australia, where Proteaceae reaches its diversity peak. Colour bar is square root transformed. Apart from a band running just inland from the west coast (which are dominated by steep escarpments), the Southwest is mostly a patchwork of remaining natural areas embedded in mostly farmland (lighter blue colours are non-natural areas).

**Supplementary Figure 5.**
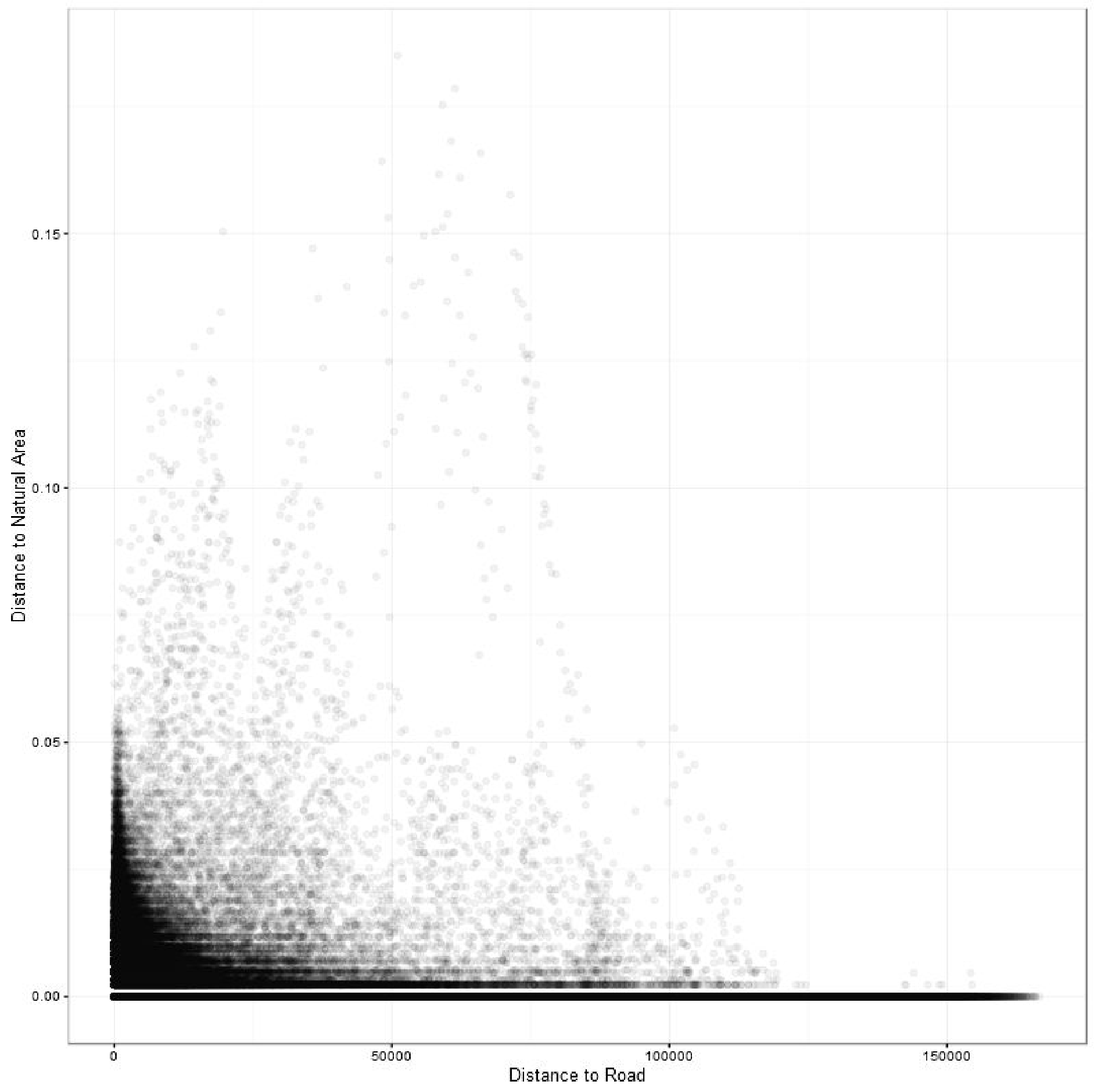
Plot showing the relationship between distance to roads and distance to natural areas for ~3 million points across Australia. There is a weak negative relationship (r = −0.11).

**Supplementary Figure 6.**
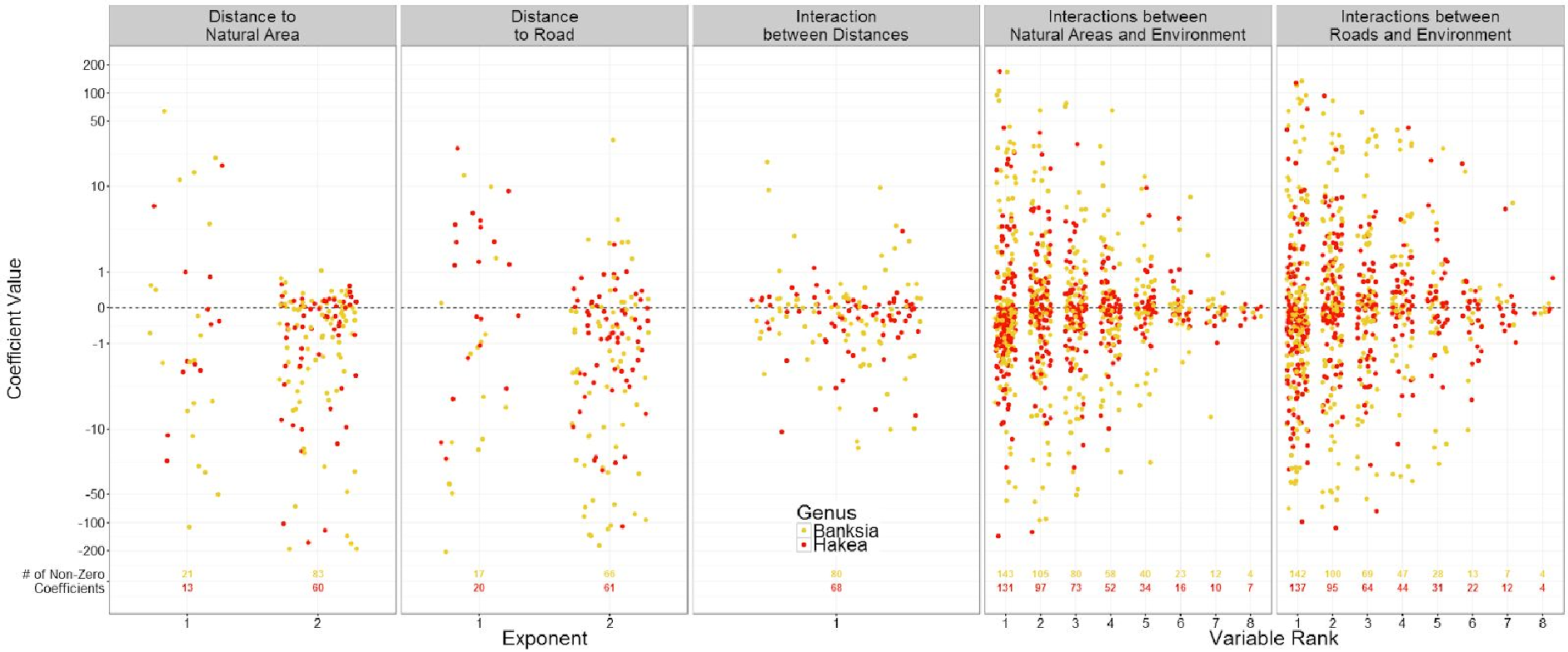
Estimated coefficients for the different effects of bias proxy variables in PPM models, for *Banksia* (yellow) and *Hakea* (red) species. The left three panels show every non-zero coefficient (those that ‘survived’ the regularization process) for distance to natural area (first panel from left), distance to road (second from left), and their interaction (middle panel). On the x axis for the first three panels is the exponent, referring to whether the coefficient was from the first or second degree polynomial expansion of the variable (second degree polynomial terms model non-linearities in the response). The right two panels show second degree polynomial coefficients involving either distance to natural areas (second from right), or distance to roads (far right panel) and environmental variables. The x axis shows the coefficients rank, ordering the coefficient values from highest absolute value (1) to lowest (8). There is a maximum of 8 second degree polynomial terms, one for each environmental variable included in the model. Below each set of coefficients is printed the number of non-zero coefficients for *Banksia* (in yellow), and *Hakea* (in red). The y axis is plotted with an arcsinh (inverse hyperbolic sin) transformation, which shrinks estimates similarly to a log transformation but allowing negative values.

